# A resurrection experiment reveals reduced adaptive potential in a common agricultural weed

**DOI:** 10.1101/2025.06.03.657543

**Authors:** Sasha G.D. Bishop, John R. Stinchcombe, Regina S. Baucom

## Abstract

**Teaser Text:** Adaptive evolution is essential for populations facing rapid anthropogenic change, yet realized evolutionary responses often fall short of expectations. Using a resurrection experiment with *Ipomoea purpurea*, we reveal that the genetic covariance structure underlying pollination and mating-system traits has become more constraining over time. Traits involved in pollinator attraction evolved within a decade, but flowering phenology—critical for coping with climate-driven shifts—was hindered by increasingly restrictive genetic covariances. Our findings suggest that temporal changes in genetic constraints can substantially dampen adaptive potential, even when selection pressures are strong.

Adaptive evolution is critical to population responses under accelerating anthropogenic global change. Although theory and some empirical work suggest that contemporary rates of environmental change can be met by equally rapid evolutionary shifts, mismatches between expected and realized evolution remain common in natural systems. Using a resurrection approach, we demonstrate that the structure of genetic covariances among pollination and mating-system traits in the common morning glory (*Ipomoea purpurea*) is increasingly constraining evolutionary trajectories. Pollinator-attracting traits evolved or became more favored on a rapid time scale of less than ten years, but genetic covariances between traits limited the adaptive rate and evolutionary trajectory of flowering phenology, a trait with widely recognized importance in adapting to climatic shifts. Our results show that changing patterns of genetic constraint can substantially reduce adaptive potential even when selection favors rapid trait evolution.

## Introduction

Understanding whether evolutionary adaptation can keep pace with rapid environmental shifts is critical to understanding current biodiversity declines and the continued resilience of ecosystems to global change (Pujol *et al*., 2018; Martin *et al*., 2023). Species can respond to rapid environmental changes by shifting their geographic range, plastic trait responses, and *in situ* adaptation. Geographic range shifts can be limited by large-scale habitat fragmentation and destruction (Lenoir *et al*., 2020; Hamann *et al*., 2021) whereas compensatory responses via plasticity have a limiting upper bound (Sgrò *et al*., 2016; Cohen *et al*., 2018). Taken together, these limits suggest that long-term, sustained responses to climatic changes rely heavily on *in situ* adaptation. A major unresolved question relating to adaptive evolution is how adaptive potential, defined here as genetic variation that can lead to a response to selection, may be changing in wild populations facing contemporary environmental pressures.

Given environmental change, a population’s adaptive capacity depends on the presence and availability of genetic variation, as well as how efficiently selection acts on phenotypic variation. Trait-trait covariances can facilitate the rate of evolutionary response if directional selection on genetically linked traits aligns with the sign of the trait covariance or slow and prohibit evolutionary response if directional selection on genetically linked traits is misaligned with the sign of the trait covariance (Etterson & Shaw, 2001; Galloway *et al*., 2018; Heblack *et al*., 2024).The presence of trait covariance in wild populations (Ćalić *et al*., 2022; Henry & Stinchcombe, 2023) as well as the possibility for these covariances to induce lags in adaptive rate in response to artificial selection (Dorn & Mitchell-Olds, 1991; Johnson *et al*., 2009; Opedal *et al*., 2022) are both well documented. However, genetic correlations can change over time in response to novel environments (Sakata *et al*., 2020; Heblack *et al*., 2024), with one of the strongest driving causes being conditions of fluctuating correlational selection (Revell, 2007; do O & Whitlock, 2023). Global change is exposing wild populations to multiple, simultaneous shifts in abiotic and biotic variables such as temperature, precipitation, salinity, extreme weather events, and shifts in population abundance or behavior of symbiotic partners.

Evidence suggests that historically coupled selective agents may be decoupling (Wadgymar *et al*., 2018), creating a context in which misaligned selection or changing correlational selection is becoming increasingly probable. This means novel correlations are reinforced by selection that maintains relationships between traits while varying trait optima, thereby providing a mechanism for sustained selection on covariance and evolution of genetic correlations. Characterizing how these genetic correlations may change in the face of global change or what agents of selection may cause changes in genetic correlations, however, remains a largely unmet challenge. As such, the causes and consequences of constraint are open questions that pertain directly to projections of adaptive rate and population viability in the face of global change.

The potential for constraints on climate change adaptation induced by among-trait genetic correlations has been assessed using a space-for-time approach (Etterson & Shaw, 2001), but spatial gradients of climatic change can be misleading proxies for climatic change over time or population response to that change (Perret *et al*., 2024). Non-climatic variables may decouple species response from climatic changes, or adaptation to local conditions can result in localized declines given climatic shifts rather than adaptation to spatially predicted trait means (DeMarche *et al*., 2021). Resurrection approaches mitigate these issues by comparing temporally sampled lineages from the same locations, thus removing spatial differences among populations altogether. While resurrection ecology is increasing in popularity for measuring phenotypic shifts, it has yet to be applied to assess the real-time evolution of constraints. Here, we combine a resurrection approach with analyses of the genetic variance–covariance structure underlying floral traits in wild morning glory (*Ipomoea purpurea*) populations and address the question of what drives the evolution of constraint by evaluating trait-specific trade-offs and pollinator visitation as a possible agent of selection.

## Materials and Methods

### Approach

We conducted a resurrection experiment using replicate maternal lines of *Ipomoea purpurea* sampled at two time points (2003 and 2012; hereafter ancestral and descendant, respectively) as seed from seven wild populations (Fig. S1). Replicates of each maternal line were planted into field conditions at the Matthaei Botanical Gardens (Ann Arbor, MI) on June 4, 2022. During the experiment we recorded trait values for five floral traits (date of first flower, corolla width, corolla length, anther-stigma distance, and nectar sucrose content (°Bx)) along with plant size (leaf count) and fitness (total seed count).

Given high genetic connectivity among *I. purpurea* populations (Alvarado-Serrano *et al*., 2019), we standardized traits within populations and pooled populations for subsequent analyses to estimate average species-level patterns rather than differences between populations (See Supplementary Materials for a population-level analysis). These measurements were used to calculate heritable genetic variation for each trait and a multivariate statistic, ***R,*** which compares the expected rate of adaptation, ***W***, given a selection gradient, **β,** and variance-covariance matrix, ***G***, to the rate of adaptation with the same selection gradient but a variance-covariance matrix with all trait covariances constrained to zero, ***Wnc*** (Agrawal & Stinchcombe, 2009). Estimates of ***R*** < 1 demonstrate evolutionary constraint such that the rate of adaptation is slower than expected without covariance. Conversely, ***R*** > 1 would indicate evolutionary facilitation. While ***G***, and consequently ***R,*** are sensitive to environmental conditions, comparisons of ***R*** in a shared environment reveal the relative difference in constraint due to genetic covariance structure between ancestral and descendant cohorts. We then investigated each trait individually by comparing predicted evolutionary change with (**Δ***z̅*) and without (**Δ***z̅***_nc_)** trait covariance to identify which floral traits influence the rate of adaptation (***R***). We further dissected temporal patterns of selection and potential genetic correlations between traits to detect instances where new or emerging pairwise trait correlations act in an antagonistic direction to selection, indicating evolution of constraints or trade-offs between traits (Roff & Fairbairn, 2012). Finally, we used pollinator visitation surveys and structural equation modeling to determine if fitness costs and benefits associated with traits are mediated through pollinator visitation frequency and thus whether adaptation in pollinator-attracting traits may constrain the response to other possible selective agents.

### Study System and Sampling Design

*Ipomoea purpurea* (Convolvulaceae) is an annual, mixed-mating vine native to the subtropical Americas and common across disturbed eastern and midwestern U.S. landscapes. Its large, showy flowers exhibit substantial natural variation in morphology and reward traits, and the species is visited by a broad assemblage of generalist pollinators (Galetto & Bernardello, 2004). These features make *I. purpurea* well suited for examining contemporary evolution in traits influencing pollination and mating system dynamics.

To assess evolutionary change over time, we conducted a resurrection experiment using seeds collected from seven wild populations in South Carolina, North Carolina, and Tennessee at two sampling points: 2003 (ancestral) and 2012 (descendant) (Fig S1). For each seed cohort, two greenhouse refresher generations were produced to reduce maternal environmental effects. Given the species’ mixed mating system (population outcrossing rates 0.2–0.8, (Kuester et al., 2017), selfed descendants remain representative of natural genotypes while enabling replicated evaluation of line means. Three replicates of each line were grown in 4-inch pots in a growth room with daily watering and a 12-hr light cycle. Plants were allowed to naturally self-pollinate, and all seeds were collected.

In May 2022, we established a common garden at the University of Michigan Matthaei Botanical Gardens. We planted 1,536 scarified seeds representing 3–8 maternal lines per population and five replicate plants per line within each of three spatial blocks (2 years x 7 populations x 3-8 maternal lines x 5 replicates x 3 blocks). Ancestral and descendant cohorts were fully randomized within blocks. Plants were staked, watered as needed, and monitored throughout the growing season. Germination exceeded 98%; however, vole herbivory reduced the final sample size to 839 individuals.

We quantified five traits associated with pollinator attraction, mating system, and phenology: corolla width, corolla length, anther–stigma distance (ASD), nectar sucrose content (°Bx, hereafter simply *nectar sucrose*), and date of first flowering (Julian date). Leaf count at flowering was recorded as an index of plant size and included as a covariate in selection models.

For each plant, four flowers sampled across the season were measured for floral morphology and nectar reward. Caliper measurements were recorded to ±0.01 mm. Nectar sucrose was estimated using standardized nectar extraction followed by refractometry. Flowers were bagged the night before sampling to prevent nectar removal. Total seed set per plant served as the estimate of fitness. Full measurement protocols are provided in SI Methods.

### Pollinator Observations

Pollinator visitation was surveyed weekly from August through early October. Observations occurred between 10:00–13:00 under suitable weather conditions and followed a standardized 20-min focal plot protocol. Across nine subplots (28–33 plants each), 277 focal plants were observed, generating 1,116 recorded visits. Pollinators were identified to order and morphospecies based on body size to maintain consistency across observers. We scored “approach” (inspection without contact) and “visit” (contact with anthers or stigma). Explicit procedures and morphogroup definitions appear in SI Methods.

### Genetic Variation and Potential for Adaptation

To determine whether traits possessed genetic variance necessary for adaptive evolution, we fit random effect models for each trait within each year:

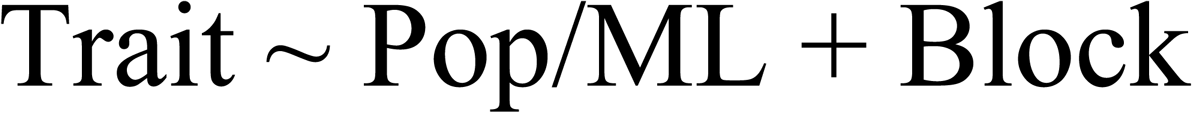

where Pop/ML represents maternal line nested within population, and Block accounts for spatial heterogeneity across the garden. Significance of the maternal-line effect was assessed via a likelihood-ratio *χ*^2^ test. Broad-sense heritability (H²) was estimated as the proportion of trait variance attributable to maternal line relative to total phenotypic variance. We repeated analyses within each population to confirm within-population variation. One population lacking variation in corolla width was excluded from subsequent analyses, as our analyses and approach require genetic variance in the focal traits. Additional model diagnostics are in SI Methods.

### Cross-Trait Constraints

To measure the impact of genetic covariances on the rate of adaptation across traits, we follow a procedure described in Agrawal and Stinchcombe, 2009. We first estimated two variance-covariance matrices using trait values standardized such that maternal line variance is unity. The first, **G,** is calculated using a multivariate mixed effects model

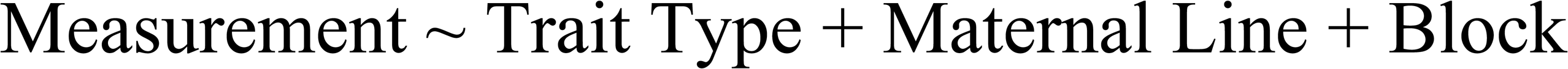

where ‘Measurement’ is a single measurement of a given trait, ‘Trait Type’ is one of the six measured phenotypic traits, and maternal line and block are both included as random effects. The second matrix, **G*_nc_***, has genetic covariances manually constrained to zero such that the diagonal of the matrix contains trait variances, but all other terms in the matrix are set to zero. Vectors of predicted response to selection, **Δ***z̅* and **Δ***z̅*_nc_ were obtained by multiplying **G** and **G**_nc_ by a vector of selection coefficients, **β f**rom a selection gradient (for information on **β** estimation see *Correlational Selection* section of SI Methods). Defining the rate of adaptation as the rate of increase of fitness of the mean phenotype, and approximating fitness by a quadratic function, we use the following equation for change in fitness of the mean phenotype (Agrawal & Stinchcombe, 2009):

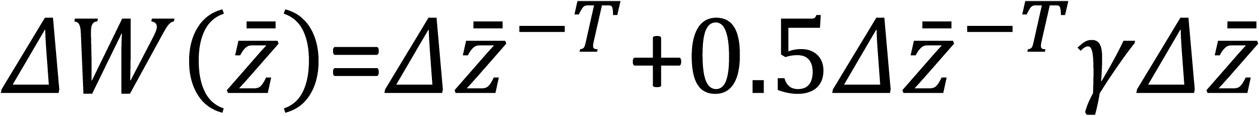

We then use 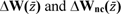 to compute a single, statistic, 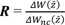 showing response to selection. This provides an intuitive metric for quantifying the rate of adaptation with covariances relative to the expected rate without them. If ***R*** = 0, then there is an absolute constraint on adaptation. If ***R*** < 1, then evolutionary constraint is present; for example, if ***R*** = 0.5 the fitness of the mean phenotype is increasing only 50% as quickly as if the traits were genetically independent. Conversely, if ***R*** > 1, then genetic covariances are facilitating evolution and accelerating the pace of adaptation.

We used bootstrap resampling to test whether the difference between evolutionary response with and without accounting for the covariance structure was significant. For each of 2000 iterations, we resampled individuals and then estimated standardized **β**, **G**, and **G_nc_** for each bootstrap sample and calculated **Δ***z̅***, Δ***z̅*_nc_, and ***R***. If 95% or more of the calculated ***R*** values were <1, we concluded that there is significant evolutionary constraint acting among traits. Each step of this analysis was performed separately for plants collected from ancestral and descendant populations. ***R_ancestral_*** and ***R_descendant_*** are then compared such that if ***R_ancestral_*** - ***R_descendant_*** > 0 for 95% or more of the resampled populations, then we conclude that genetic covariances constrain evolution less in ancestral populations.

A comparison of ***R*** across years provides a metric for changes in how strongly evolutionary constraints impact adaptive rate (*i.e.* the discrepancy in response to selection with and without covariance), however it does not directly measure whether the absolute adaptive rate changes between ancestral and descendant populations. To do this, we implement a modified ***R*** statistic, ***R’*** where 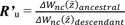 represents a predicted difference in response between ancestral and descendant populations to selection without constraints (*i.e.* a **u**nivariate analysis of how adaptive potential changes over time).

Similarly, 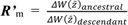 represents a predicted difference in response between ancestral and descendant populations to selection with constraints (*i.e.* a **m**ultivariate analysis of how adaptive potential changes over time). If ***R’*** < 1 for 95% or more of the resampled populations, we conclude that the adaptive rate is lower in ancestral populations, meaning that adaptive capacity is sustained or even increasing over time. Conversely, if ***R’*** > 1 for 95% or more of the resampled populations, we conclude that the adaptive rate is greater in ancestral populations, meaning that adaptive capacity is decreasing over time.

### Trait-Specific Constraints

The existence of evolutionary constraint across a suite of traits does not guarantee every trait is impacted in the same manner. To assess which traits underlie changes in ***R*** and ***R’*,** we compare **Δ***z̅*_ancestral_ to **Δ***z̅*_descendant_ for each trait and **Δ***z̅* to **Δ***z̅*_nc_ for each trait and year independently. If 95% or more values of **Δ**z*_i_* from the 2000 bootstrap samples described above are less than ***Δ****z_i_nc_*, we conclude that evolution in that trait in that year is subject to adaptive constraint. If there is no significant difference between **Δ**z*_i_* and **Δ**z*_i_nc_*, then there is no constraint or facilitation on that trait, and if **Δ**z*_i_* > ***Δ****z_i_nc_*, then the trait is subject to adaptive facilitation. If 95% or more values of **Δ***z̅*_descendant_ < **Δ***z̅*_ancestral_, then we conclude that predicted adaptive rate with covariance has decreased over time, whereas if **Δ***z̅*_descendant_ > **Δ***z̅*_ancestral_, then predicted adaptive rate has increased and no difference between the two indicates a stable adaptive rate.

### Trait Correlations & Conflicting Selection

We quantified Pearson correlations among maternal-line means for all trait pairs and tested for temporal shifts in correlations via 10,000 bootstrap replicates. Directional agreement or conflict between direct selection (**β**) and trait correlations was evaluated to determine whether emerging trade-offs aligned with observed covariance evolution.

To assess total selection on traits as a metric for adaptive capacity, we estimated selection differentials (*S*) from univariate regressions of relative fitness on each trait, with traits standardized to variance of one and block as a random effect. Total stabilizing or disruptive selection were assessed with a near-identical model including both the linear and quadratic terms for a given trait. Selection gradients (β, γ) were obtained from a multivariate regression including all traits simultaneously. Traits were standardized such that each maternal-line variance equaled one. Significance was assessed using Type III ANOVA. Extended correlation tests and selection models are provided in SI Methods.

### Pollinators as Agents of Selection: Structural Equation Models

To test whether evolution may be mediated by pollinator behavior, we evaluated seven *a priori* structural equation models representing alternative causal structures linking floral traits, pollinator visitation (approach and visit), and fitness in the ancestral populations. The hypotheses tested include five nested hypotheses regarding the extent to which pollinators mediate the impact of floral traits on fitness, and two non-nested hypotheses - one testing whether relationships are correlated but not causal and another testing whether all trait variation is determined by plant size. Analyses were conducted using piecewiseSEM in R, which accommodates mixed-model components and non-Gaussian data types such as count (R Core Team, 2024; Lefcheck, 2016).

Model fit was assessed using a log-likelihood **χ²** statistic comparing model-implied and observed variance-covariance relationships; non-significant (p > 0.05) **χ²** indicates adequate fit. One model lacked degrees of freedom and was assessed via AIC comparisons. Path coefficients and indirect effects were estimated for only the best-fit model and statistical power was assessed using pwrSEM (Wang & Rhemtulla, 2021). All *a priori* hypotheses, model structures and fit statistics are included in SI Methods and Tables S4–S7.

### Realized Phenotypic Evolution

To evaluate phenotypic change between years, we fit mixed models with year, population, and their interaction as fixed effects and block as a random effect:

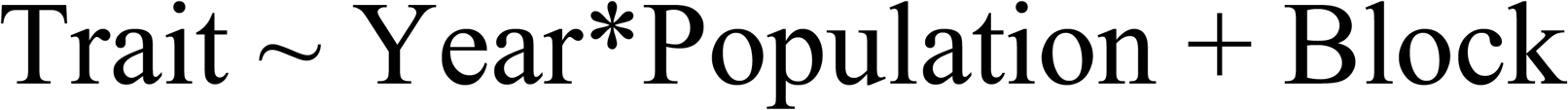

Traits were log-transformed to improve normality. Post hoc comparisons of estimated marginal means were conducted using the emmeans package in *R* (R Core Team, 2024; Lenth, 2025). Full model outputs appear in SI Methods.

## Results

### Trait Covariance is Causing a Rapid Decline in Adaptive Rate Over Time

In both the ancestral and descendant populations, across traits, the overall rate of adaptation is lower when genetic covariances between traits are considered. Notably, however, evolutionary constraint is much stronger in the descendant populations with ***R*** = 0.09 compared to ***R*** = 0.76 in the ancestral populations (99.85% significance threshold; Fig. 1A). Thus, in the ancestral populations, accounting for trait covariances results in an adaptive rate that is 76% as fast as predictions without trait covariances, but this predicted rate drops precipitously to 9% in the descendant populations, indicating that constraints have increased over time. An alteration to the ***R*** statistic, ***R’*** (with values >1 indicating greater adaptive potential in ancestral populations), that compares the ancestral and descendant populations directly (see SI Methods) shows that predicted rates of adaptation without covariance are largely sustained over time, though perhaps marginally faster in ancestral populations, ***R_nc_’*** = 1.81. However, when accounting for trait covariance, ***R’*** = 23.61, demonstrating that, contrary to univariate predictions, the adaptive rate was 23x faster in ancestral populations - *i.e.* over nine years the populations lost 96% of their adaptive potential. These findings indicate that constraints among floral traits evolve on a rapid time scale of less than ten years and are acting to limit adaptive potential, overall highlighting that consideration of multi-trait dynamics is critical to understanding the impact of global change on patterns of biodiversity.

**Figure 1.**
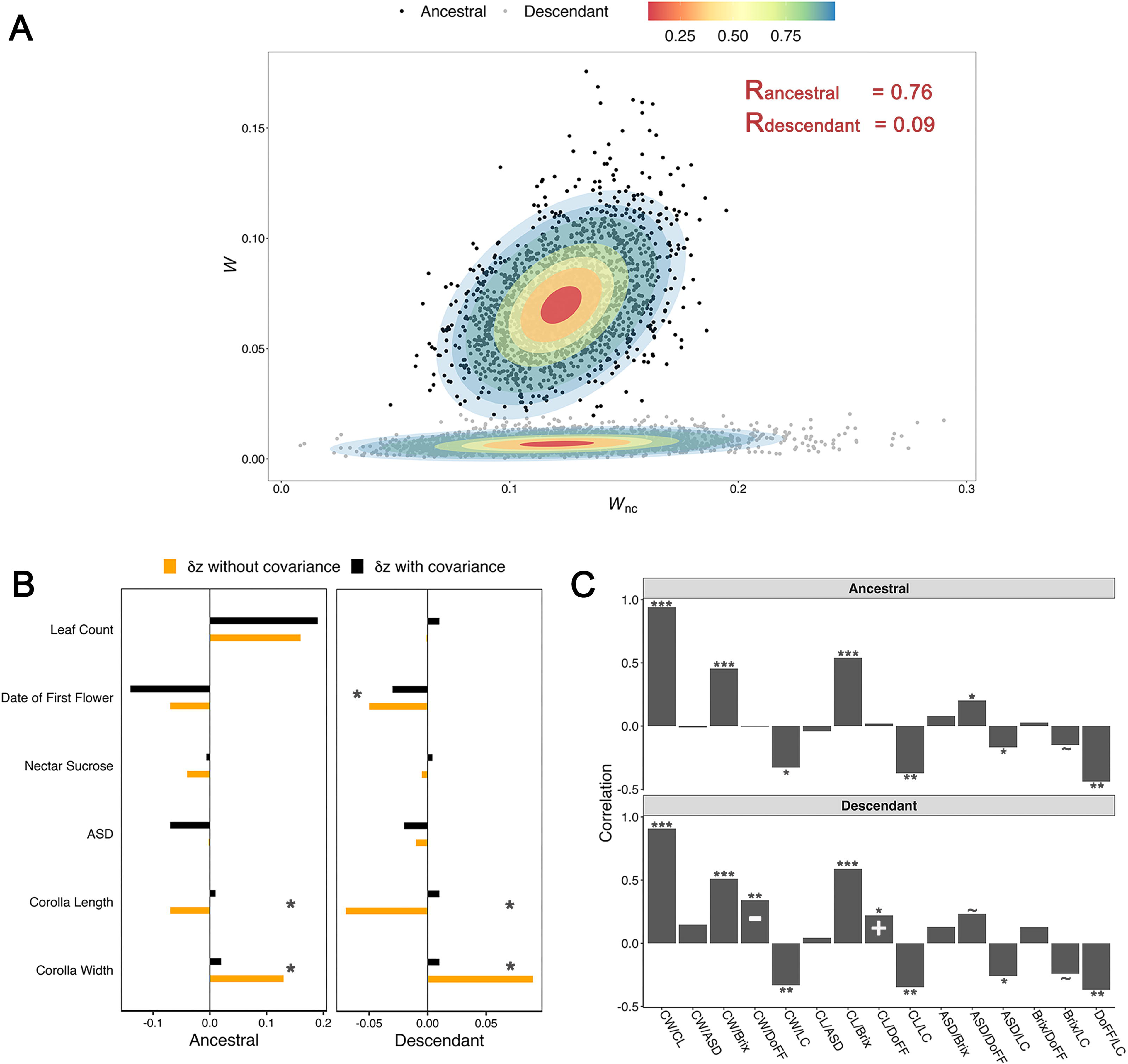
Adaptive potential decreases over time due to a trade-off between corolla size and flowering phenology. **(A)** Bootstrap estimates of ***R***, plotted as the two component parts of the ratio ΔW(*z̅*) and ΔW_nc_(*z̅*). Black points indicate resampled ***R*** estimates for ancestral populations, and grey points indicate resampled ***R*** estimates for descendant populations. Ellipses correspond to confidence intervals of 10, 34, 50, 68, 90, 95, and 98% with the respective color gradient ranging from red to light blue. Actual ***R*** estimates are reported in the top right. (**B)** The expected response to selection with (Δz_i_) and without (Δz_i_*nc*_) trait covariances (black and orange bars, respectively) for each trait. Traits shown with an asterisk had a significant difference between Δz*_i_* and Δz*_i_nc_* in either ancestral or descendant populations, based on bootstrap resampling. **C)** Pairwise correlations between traits for both ancestral and descendant populations calculated as Pearson’s correlation coefficient. Trait labels are as follows: CW = Corolla Width, CL = Corolla Length, ASD = Anther-Stigma Distance, Brix = Nectar Sucrose, DoFF = Date of First Flower, LC = Leaf Count. Traits displaying a significant change in correlation over time based on bootstrap resampling are shown with a “-” indicating that the direction of joint selection is antagonistic to the direction of correlation or a “+” indicating that the direction of selection is reinforcing to the direction of trait correlation. Asterisks next to each bar indicate a significant correlation within year (p < 0.1∼, p < 0.05*, p < 0.01**, and p < 0.001***). **Alt Text:** Graphical representations of a comparison of ancestral and descendant populations for three separate statistcs: ***R,*** Δz with and without covariances, and trait correlations. Sub-panels highlighting each of those statistcs are labeled A-C.

We next compared expected adaptation of individual traits with and without covariance (**Δ***z̅* to **Δ***z̅*_nc_) to identify which traits underlie observed shifts in ***R*.** We found significant (>95% significance threshold) constraint on the rate of adaptation in corolla width as well as a change in the direction of evolution for corolla length in both the ancestral and descendant populations (corolla width, ancestral: **Δ**z_cw_ = 0.02, **Δ**z_nc_cw_ = 0.13, descendent: **Δ**z_cw_ = 0.01, **Δ**z_nc_cw_ = 0.09; corolla length, ancestral: **Δ**z_cl_ = 0.01, **Δ**z_nc_cl_ = - 0.07, descendent: **Δ**z_cl_ = 0.01, **Δ**z_nc_cl_ = −0.07; Table 1, Fig. 1B). Comparatively, for anther-stigma distance (ASD) and day of first flowering, we find that genetic covariance between traits in ancestral populations results in adaptive facilitation such that expected adaptation of smaller ASD (**Δ**z_ASD_ = −0.07, **Δ**z_nc_ASD_ = −0.002; 96.55% significance threshold) and earlier flowering (**Δ**z_DoFF_ = −0.14, **Δ**z_nc_DoFF_ = −0.07; 99% significance threshold) is faster than without covariance. Descendant populations, however, display no effect of covariances on ASD, and show adaptive constraint on flowering phenology such that the adaptive rate of flowering time is slowed by covariance with other traits (**Δ**z_DoFF_ = −0.03, **Δ**z_nc_DoFF_ = −0.05; 92% significance threshold; Table 1, Fig. 1B). A comparison of **Δ**z between years shows that there is a decrease over time in expected rate of adaptation toward earlier flowering (Table 1, 99.85% significance threshold) and plant size (ancestral **Δ**z_size_ = 0.19, descendant **Δ**z_size_ = 0.01; 99.95% significance threshold).

**Table 1.** The adaptive potentials of floral traits are limited by trait-trait covariances. For each trait, the mean and standard error for Δz*_i_* and Δz*_i_nc_*, with those displaying significant evolutionary constraint marked in bold. Significance is defined as the percent of bootstrap samples where Δz*_i_* - Δz*_i_nc_* is either above or below zero, depending on the expected difference calculated from the actual data. Δz is calculated with standardized traits such that variance(maternal line) = 1. Significant broad sense heritabilities (*H^2^*) for each trait are also shown in bold. Estimated marginal mean trait values are shown for corolla width, length and ASD in cm, nectar sucrose in °Bx, flowering phenology as Julian Date, plant size as leaf count, and relative fitness as seed count. Ancestral and descendent populations are noted as A or D, respectively. Bolded values represent traits with a significant change in trait value between ancestral and descendant populations (Table S1).

| | Year | $\Delta z_i$ | SE | $\Delta z_{i\_nc}$ | SE | $H^2$ | Trait Mean |
| --- | --- | --- | --- | --- | --- | --- | --- |
| **Corolla Width**<br>(mm) | A | **0.02** | **1.86E-05** | **0.13** | **2.23E-05** | **0.11** | **48.0** |
|  | D | **0.01** | **5.37E-06** | **0.09** | **1.54E-05** | **0.10** | **48.7** |
| **Corolla Length**<br>(mm) | A | **0.01** | **1.70E-05** | **-0.07** | **1.36E-05** | **0.09** | 53.4 |
|  | D | **0.01** | **4.50E-06** | **-0.07** | **1.12E-05** | **0.05** | 53.4 |
| **ASD**<br>(mm) | A | -0.07 | 2.11E-05 | -0.002 | 3.16E-07 | **0.08** | 0.37 |
|  | D | -0.02 | 7.14E-06 | -0.01 | 2.13E-06 | **0.09** | 0.55 |
| **Nectar Sucrose**<br>(°Bx) | A | -0.006 | 1.88E-05 | -0.04 | 8.77E-06 | **0.10** | 3 |
|  | D | 0.004 | 4.76E-06 | -0.005 | 1.07E-06 | **0.10** | 3 |
| **Flowering Phenology**<br>(Julian Date) | A | -0.14 | 1.91E-05 | -0.07 | 8.33E-06 | **0.30** | **218** |
|  | D | **-0.03** | **7.11E-06** | **-0.05** | **4.02E-06** | **0.13** | **217.3** |
| **Plant Size**<br>(Leaf Count) | A | 0.19 | 3.32E-05 | 0.16 | 3.01E-05 | **0.09** | **37.2** |
|  | D | 0.01 | 6.05E-06 | -0.001 | 1.24E-07 | **0.03** | **33** |

### Constraints Result in New Trait Optima

Two commonly proposed mechanisms for lack of evolutionary change are an absence of heritable genetic variation or trade-offs in the form of negative genetic correlations or covariances (Heblack *et al*., 2024); either, or both, of these mechanisms could explain the results on adaptive rate. However, we found each of the floral traits to exhibit heritable genetic variation across collection years (Table 1). This means that lack of appreciable genetic variation within individual traits will not induce evolutionary constraint, further corroborated by variation maintained in ***W_nc_*** but not in ***W*** (Fig. 1A). Analysis of genetic correlations showed that, while trade-offs between traits were present and generally took the form of negative genetic correlations between floral traits and plant size, most remained similar in sign and strength between the ancestral and descendent populations (Fig. 1C, S2). Two notable exceptions are the correlation between corolla width or corolla length and date of first flower (corolla width and date of first flower: ancestral r = −0.005, p = 0.481, descendant r = 0.34, p < 0.0001; corolla length and date of first flower: ancestral r = 0.02, p = 0.627, descendant r = 0.22, p = 0.003). The emergence of a positive correlation between corolla size and flowering date in descendant populations indicates that a new trade-off has evolved between larger flowers and earlier flowering.

We likewise identified different trait optima in the descendent versus ancestral populations favoring evolutionary investment in pollinator-attracting traits at the cost of evolvability in flowering phenology. In the common garden setting, where the selection regime is shared, a difference in selection coefficient between the ancestral and descendent populations means that there is a different fitness response for a given trait in that shared environment. As such, we report comparisons of total and direct selection on traits to show shifts in adaptive capacity.

We found that total selection on the date of first flower differed between ancestral and descendant populations (ANCOVA contrasting slopes: F = 5.39, p = 0.020, Table S2). In ancestral populations, we uncovered higher fitness in earlier flowering plants (S = −0.20, p = 0.02; Table S2) as well as a marginally-significant phenotypic shift to earlier flowering in descendant populations by ∼1 day (t-ratio = 1.78, p = 0.07; Fig. S3, Table S1, *also shown as a significant shift in greenhouse conditions*, (Bishop *et al*., 2023)). In contrast, in line with the constraint indicated by Δz_DoFF_ and a trade-off with flower size, earlier flowering dates are no longer favored in descendant populations, and instead highest fitness is found for intermediate flowering dates (γ = −0.12, p = 0.002; Table S2). Descendent populations also displayed the emergence of marginally-significant, but novel correlational selection between date of first flower and three other traits: corolla width (γ = −0.34, p = 0.071), nectar sucrose content (γ = −0.14, p = 0.067), and corolla length (γ = 0.39, p = 0.045, Fig. 2).

**Figure 2.**
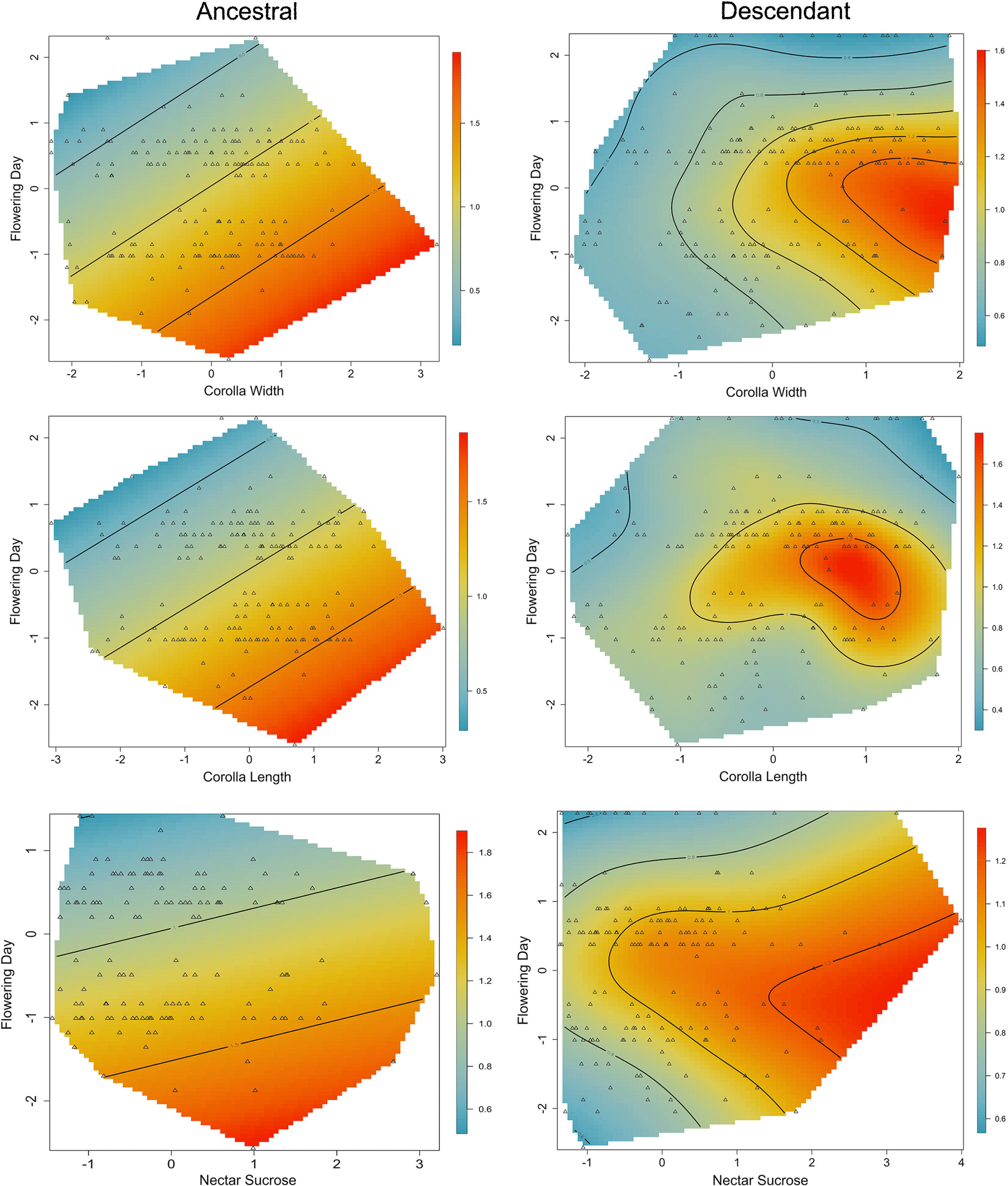
Correlational selection is present in descendent but not ancestral populations. Fitness surface for correlational selection in ancestral and descendant populations acting upon date of first flower and three other floral traits tested using selection gradients including all linear, non-linear, and interacting terms (multivariate selection). Relative fitness is depicted by the color gradient with red being the highest fitness, yellow intermediate fitness, and blue lowest fitness. Triangles represent the measurements underlying the model. In ancestral populations, no indication of correlational selection was present. All traits were included in the selection gradient, only those with selection gradients indicative of correlational selection in descendant populations are shown: corolla width (γ = −0.34, p = 0.071), corolla length (γ = 0.39, p = 0.045), and nectar sucrose (γ = −0.14, p = 0.067). **Alt Text:** Six panels separated into two columns (ancestral and descendant populations) with each panel displaying a fitness surface for correlational selection between flowering time and another floral trait.

Despite constraints on the rate of adaptation of corolla width, both corolla width and nectar sucrose display indications of increasing investment in pollinator-attracting traits. Specifically, corolla width shows significant positive total selection in both the ancestral and descendent populations (S = 0.14, p < 0.001 ancestral; S = 0.14, p < 0.001 descendant; Table S2); further, the continued evolution of broader corollas is corroborated by a significant increase in mean trait value from ancestral to descendant populations by ∼1mm (t-ratio = −2.53, p = 0.01; Fig S4, Table S1). Although total selection indicates positive selection for corolla length (S = 0.11, p = 0.002 ancestral; S = 0.06, p = 0.031 descendant), analysis of direct selection shows no selection in ancestral populations and negative selection on corolla length in descendant populations (β = −0.37, p = 0.03; Table S3). Thus, the positive total selection on corolla length likely arises from the strong correlation between corolla length and width (r = 0.92, p < 0.001) and the overall positive selection on corolla width. Evidence for nectar sucrose is weaker than with corolla width: ancestral populations display a pattern of disruptive total selection where both high and low levels of sucrose are favored (*γ* =0.07, p = 0.02; Table S2), however, there is a marginally-significant change in the quadratic relationship between fitness and nectar sucrose in descendant populations (ANCOVA contrasting slopes: F = 3.40, p = 0.066, Table S2) such that lower sucrose values are no longer favored and instead there is selection for high values of nectar sucrose (*S* = 0.07, p = 0.04, Table S2). However, there is not direct selection in descendant populations or a significant shift in mean trait value, such that, as with corolla length, this pattern may partially reflect covariance with other floral traits. Nonetheless, the change in quadratic relationship as well as structural equation modeling indicates that nectar sucrose contributes indirectly to plant fitness through pollinator attraction (see *Pollinators as Agents of Selection,* below), suggesting a functional role in pollinator-mediated selection despite limited evidence for contemporary evolutionary change.

Overall, these data suggest that the evolution of flowering phenology in descendant populations appears increasingly constrained by covariance with floral attraction traits, particularly corolla width, alongside the emergence of correlational selection that favors intermediate flowering dates conditional on floral phenotype (Fig. 1B & 1C, Tables S2 & S3). These changes are additionally accompanied by a decrease in relative fitness of descendant populations compared to ancestral (ancestral = 1.06, descendant = 0.93; p = 0.001), indicating that, at least in some environments, reduced adaptive capacity is also resulting in lowering fitness. Beyond the rapid evolution of trade-offs, our results show that constraints due to changes in genetic architecture have evolved within the nine-year period studied, and that the covariance structure—particularly between floral traits and the timing of flowering—may remain unstable as global changes continue.

### Pollinators as Agents of Selection

We next examined the potential that pollinators impose selection on both corolla width and nectar sucrose content in *I. purpurea* using structural equation modeling. Our best supported model (Table S4, S5) indicates that increases in corolla width and nectar sucrose content between ancestral and descendant populations are driven by pollinator preference, whereas changes in flowering phenology are not responsive to pollinator foraging frequency, but rather some other environmental factor. Specifically, we find a positive effect of corolla width on frequency of pollinator floral approaches (r = 0.250, p = 0.02) and pollinator visits (r = 0.337, p = 0.001) as well as a positive relationship between nectar sucrose content and pollinator approaches (r = 0.148, p = 0.002), meaning that large corolla widths and high sucrose content in nectar attracts more pollinators to a flower (Fig. 3, Table S6). An additional high conversion of pollinator approach to foraging (r = 0.373, p < 0.0001) and positive relationship between pollinator foraging and plant fitness (r = 0.171, p = 0.033) results in an indirect relationship between corolla width and nectar on plant fitness where high values of each are favored by pollinators and result in higher plant fitness. Conversely, earlier dates of first flower have a positive effect on plant fitness (r = 0.325, p = 0.004), but do not impact pollinator foraging behavior, meaning that selection on flowering phenology is driven by something other than pollinators, an observation corroborated by numerous studies showing that flowering phenology is core to adaptation to global change factors such as temperature (Dai *et al*., 2017; Cheptou *et al*., 2022) and precipitation (Chand *et al*., 2022). Overall, and in conjunction with the development of constraints on flowering phenology due to trade-offs with corolla width and first flower, our results highlight an adaptive pathway that is driven by investment in pollinator attraction at the cost of evolvability in flowering phenology, a shift that has occurred within a nine-year period.

**Figure 3.**
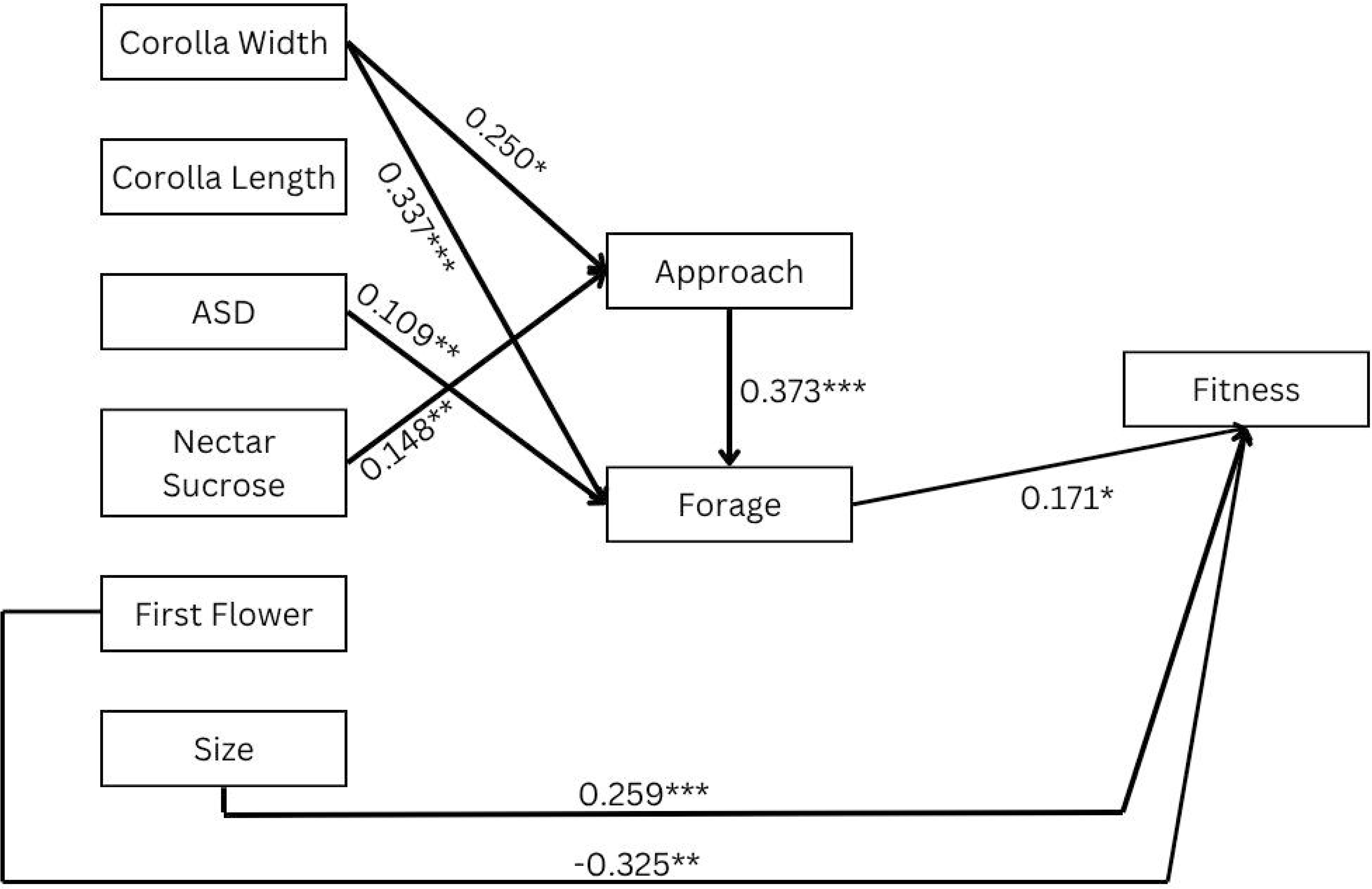
Pollinator foraging behavior mediates fitness effects of floral size and quality traits. Best-fit SEM showing the direct and indirect effects determining pollinator visitation rate and fitness. Standardized path coefficients are shown for each significant path where * indicates a p-value < 0.05, ** indicates p < 0.01, and *** indicates p < 0.001. See Table S6 for a full listing of direct, indirect, and total effect sizes of functional traits on fitness. **Alt Text:** Path diagram showing the direct and indirect relationships between floral traits (on left), pollinator approach and foraging (middle), and plant fitness (on right).

## Discussion

A central question in evolutionary biology is whether adaptive evolution can keep pace with rapid, multi-dimensional environmental change. Although models often predict rapid evolutionary responses, empirical studies repeatedly uncover lags between expected and realized adaptation in the wild (Martin *et al*., 2023). Using a resurrection framework, we demonstrate that genetic constraints among floral traits in *Ipomoea purpurea* have strengthened markedly over just nine years, resulting in a 96% reduction in predicted adaptive rate in the common garden setting. These results provide a mechanistic explanation for why adaptation may stall even in populations with abundant genetic variation: rapid evolution of trait covariances can fundamentally reorganize the genetic architecture that determines how efficiently selection translates into evolutionary change.

Our findings reveal that constraints themselves evolve and do so on ecologically relevant timescales. Ancestral populations showed moderate constraint, whereas descendant populations exhibited strong antagonistic covariances—particularly involving corolla width and flowering phenology—that greatly reduced predicted evolutionary response. Importantly, heritable variation was present for all focal traits, indicating that limits to adaptation arise not from the loss of genetic variance, but from the reorganization of variance across traits. Strengthened correlations between floral size and flowering time in descendant populations mark the evolution of a new trade-off linking investment in pollinator attraction to delays in flowering, thereby shifting multivariate adaptive trajectories.

Pollinators appear to be central to this shift. Structural equation models show that larger corollas and higher nectar sucrose content increased pollinator visits and, indirectly, plant fitness, whereas pollinators did not mediate the fitness effects of flowering time. This asymmetric influence means that selection by pollinators pushes floral size upward but simultaneously creates antagonistic covariance with flowering phenology. In effect, adaptation toward greater pollinator attraction reshapes the covariance structure in ways that constrain the evolution of earlier flowering, a trait strongly tied to climatic adaptation in many plant species. These findings align with emerging evidence that global change factors may not act independently, but instead impose conflicting (Wadgymar *et al*., 2018) or interacting (García *et al*., 2023) selective pressures that channel adaptation in opposing directions.

Several considerations provide context for these inferences. First, distinguishing drift from selection as the mechanism behind changes in covariance structure is challenging. However, if drift were the major agent underlying the changes in covariance structure that we document, we would expect to see proportional change in variances and covariances across traits (Chapuis *et al*., 2008; Doroszuk *et al*., 2008). However, we observed changes in genetic correlations that were uneven across the traits (Fig. 1C) and that the evenness (*E_ƛ_*) of **G** declines from ancestral (*E_ƛ ancestral_* = 0.62) to descendant (*E_ƛ descendant_* = 0.49) populations. Together, this indicates that genetic variance becomes less evenly distributed and that drift is an unlikely mechanism (Agrawal & Stinchcombe, 2009). Second, we used selfed maternal lines and a single common garden experiment, which allowed us to reduce genetic heterogeneity and more directly attribute differences between sampling years to evolutionary rather than mating-system variation. We note that this approach may not fully capture covariance structure as expressed in naturally outcrossed populations; however, given that *I. purpurea* freely self-pollinates across much of its range (Kuester *et al*., 2017), selfed lines offer a biologically appropriate and experimentally tractable means of minimizing within-line variation and isolating temporal genetic differences. Given that *I. purpurea* is capable of autonomous selfing, we also acknowledge that pollinator-mediated selection in this system may act through variation in pollen export and mating dynamics rather than pollen receipt alone. Although the absolute **G-**matrix values will be environmentally sensitive, the single common garden thus allows for a direct comparison between time points of direction and relative magnitude of evolutionary constraint. Finally, previous work indicates high genetic connectivity among *I. purpurea* populations (Alvarado-Serrano *et al*., 2019), so we standardized traits within populations and analyzed them collectively to estimate species-wide responses to an average pattern of selection rather than focus on population-specific differences. As a diagnostic, we also calculated ***R*** and **Δ***z̅* for each population (Table S9), but models fitted separately had limited power to estimate higher-order interaction terms. Pooling populations therefore provides a more robust estimate of the typical strength and sign of these interactions. Ultimately, these caveats do not diminish the core conclusion: the covariance structure underlying floral traits has evolved rapidly, with major consequences for adaptive potential.

Prior work has emphasized shifts toward increased selfing (*Viola praemorsa*, Jones *et al*., 2013; *Viola arvensis*, Cheptou *et al*., 2022), investment in pollinator attraction due to reduced pollinator abundance (*Lobelia siphilitica,* Brown & Caruso, 2023), or climatically-induced shifts in flowering phenology (Freimuth *et al*., 2022). Together, our results identify another pathway by which plants may respond to global change: the evolution of genetic constraints themselves. We show that selection favoring pollinator-attracting traits may indirectly restrict the evolution of flowering time, a trait critical for climatic adaptation. This has important implications for population persistence.

Recent decades have seen significant reductions in pollinator populations (Winfree *et al*., 2011; Thomann *et al*., 2013; Hallmann *et al*., 2017), with unprecedentedly high declines of bumblebee species in North America specifically between the years of 2000-2014 (Soroye *et al*., 2020) when these experimental *Ipomoea purpurea* populations were collected. Furthermore, the southeastern U.S. where these plants were collected has witnessed climatic changes such as rising temperatures and an increase in drought conditions interspersed with extreme rainfall events (Jay et al., 2018). While the evolution of flowering time is well established as a key mechanism for flowering plants to adapt to climatic changes in temperature and precipitation (Byers, 2017; Renner & Zohner, 2018; Gérard *et al*., 2020; Soares *et al*., 2021), our research suggests that declining pollinator populations could undermine resilience to climate change through adaptive constraint–specifically, the diminished adaptive potential in flowering phenology is at odds with projected climatic shifts that continue to alter temperature and precipitation patterns (Pörtner *et al*., 2022). These findings thus highlight the necessity of considering multiple dimensions of global change, and emphasize that adaptation to one factor, such as reduced pollinator populations, could conflict with the adaptive responses required for other factors, like climate-induced shifts in temperature and precipitation.

## Acknowledgments

The authors thank Michael Palmer and Jeremy Moghtader at the Matthaei Botanical Gardens for assistance with experiment logistics and field preparation. We also thank Grace Zhang and the team of undergraduate field assistants who helped with field maintenance and trait measurements. We give particular thanks to field technicians Anah Soble, Alice Eliott, and Ruby Barron for assistance with data collection and pollinator observations, and to Adam Kuester for the 2012 seed collections. Management of plants in a second growth room resurrection experiment used to look at plasticity was done primarily by Lily MacKrell and Alex Husted. Finally, we thank Drs. Rachel Spigler, Timothy James, and Meha Jain for their invaluable feedback on experimental design and analysis.

## Author Contributions

S.B. and R.S.B. conceived and designed the study with R.S.B. acting as advisor. S.B. collected and analyzed the data and J.R.S. provided critical review and analysis recommendations. S.B. and R.S.B drafted the initial manuscript with all authors contributing to the final version.

## Data and materials availability

Primary data and R code used in these analyses are made available through the following digital dryad repository: https://datadryad.org/dataset/doi:10.5061/dryad.2z34tmpz8

