## Supplemental methods, figures, and tables for "A resurrection experiment reveals reduced adaptive potential in a common agricultural weed"

**This PDF file includes:**

Supporting text: Materials & Methods
Figures S1 to S4
Tables S1 to S9
SI References

**Supporting Information Text**

**Materials & Methods**

**Trait measurements**–We recorded leaf count as a metric for plant size and date of flowering onset for each plant and measured floral morphology and reward traits for four flowers per plant. Corolla width, corolla length, and anther-stigma distance were measured using digital calipers (precision: ±0.01mm), with anther-stigma distance representing the difference in height between the stigma and the tallest anther. To extract nectar from the flower, 10uL of reverse osmosis (RO) water was pipetted directly into the base of a flower, pushing the pipette tip past the base of the stamens and pipetting up and down to mix and extract nectar. We then quantified sucrose content of this nectar/water solution using a pocket refractometer to record percent mass sucrose (°Brix, hereafter nectar sucrose content) in flowers protected with a mesh cover the night before to control for nectar removal. Finally, we collected and counted the number of seeds produced by each plant and used the total seed set as our estimate of fitness.

**Pollinator observations–**Pollinator observations were conducted once a week during the months of August, September, and early October, always between the hours of 10AM – 1PM. In total, a field team observed pollinator visitation of 277 plants distributed across nine sub-plots, each with 28-33 plants. Each observation round had a duration of twenty minutes, and eight rounds per plot were conducted in total resulting in 1,116 observed pollinator visits. Each week, we rotated the plot observed at 10AM to balance the influence of time of day across plots. To standardize across different observers and varying levels of insect taxonomic expertise, we identified pollinators to the level of order and then morphospecies based on size. The types of pollinators we looked for were: large (10-15mm) and extra-large (>15mm) social bees in the family Apidae; small (<5mm) and medium (5-10mm) solitary bees in the families Halictidae or Apidae, large (>10mm) solitary bees in the families Andrenidae or Megachilidae; bee flies (family Bombyliidae), small (<10mm) and large (>10mm) syrphid flies in the family Syrphidae; other flies (families Muscidae and Calliphoridae), Lepidoptera, Coleoptera, Orthoptera, Hemiptera, wasps (family Vespidae), ants, and other. We recorded pollinator approach as a pollinator flying up to a flower, but not contacting the stigma or anthers. Pollinator foraging was recorded as a pollinator directly contacting the anthers and/or stigma.

**Genetic variation and potential for adaptation**–We determined if there was evidence of genetic variation for each floral trait by first looking at whether maternal line has a significant effect on phenotypic trait variation using a likelihood ratio $\chi^2$ test using the lmerTest package in R (Bates *et al.*, 2015). We also calculated broad sense heritability for each trait in ancestral and descendent populations as the proportion of variance in the model attributable to maternal line (i.e. $V_G / V_G + V_{Block} + V_{Residual}$). We repeated this set of analyses for each population individually to affirm that there is sufficient genetic variation not just across, but also within populations for adaptation to occur. A single population (population 12, Fig. S1) lacked variation in corolla width, and was removed from subsequent analysis. A diagnostic check of cross-population analyses including population 12 displays lower significance, but the same directional pattern of change over time (Table S8).

**Trait correlations**–We first use the cor.test function in base R ("The R Project for Statistical Computing," 2024) to calculate Pearson's correlation coefficients between maternal line means for traits and the t-statistic of each pairwise correlation to test significance. Differences in correlation between years is evaluated by bootstrap resampling trait values and calculating Pearson's correlation coefficient 10,000 times. If the difference between the Pearson's correlation coefficient in ancestral and descendant populations is greater than zero >95% of the time, we conclude that two traits have significantly decreased in correlation over time. If the difference is less than zero >95% of the time, we conclude that correlation between two traits has significantly increased over time.

To determine whether changes over time in trait correlations are indicative of the emergence of new trade-offs in the common garden selective environment, we tested whether the direction of selection on pairs of traits (*i.e.* $\boldsymbol{\beta}_{\text{Trait 1}}$ x $\boldsymbol{\beta}_{\text{Trait 2}}$) opposes the sign of the trait correlation (Etterson & Shaw, 2001; Galloway *et al.*, 2018).

**Correlational selection**–To assess whether there are additional indications of new trait optima or if correlational selection could be a driving force behind changes in covariance structure and trait correlations, coefficients of cross-product terms were extracted from a multivariate regression model. For this model, focal traits corolla width, corolla length, anther-stigma distance, nectar sucrose content, date of first flower, and leaf count were all standardized to a mean of zero and standard deviation of one across all data, and relative fitness was measured as the total seed set for an individual plant divided by the mean seed count for that year. Leaf count is included in initial model testing as a plant size covariate to ensure that fitness effects are not just due to plant size. However, due to model saturation and lack of significance of terms including leaf count, we removed it from the final model. The model follows the general form of relative fitness regressed on all phenotypic traits together including linear terms ($\boldsymbol{\beta}$), quadratic terms, and cross-product terms of all focal traits; quadratic estimates and standard errors for focal traits, but not cross-products between traits, were doubled to estimate $\boldsymbol{\gamma}$. $\boldsymbol{\beta}$ calculated in this manner is also used for the $\boldsymbol{\Delta}$z and $\boldsymbol{R}$ calculations.

The significance of interactive effects between traits were evaluated separately for ancestors and descendants, where significance indicates that a combination of focal traits maximizes fitness. We visualized fitness surfaces for significant cross-product terms by performing a thin-plate spline nonparametric regression approach, using the Tps function in the R fields package (Nychka *et al.*, 2015). Smoothing parameters for each spline were chosen to minimize generalized cross-validation score.

**Total selection on traits**–To look directly at whether traits found to be involved in trait trade-offs show declines in adaptive capacity (*i.e.* ability to respond to the same selective pressure) in line with predicted decreases in adaptive rate seen in $\boldsymbol{\Delta}$z, we assess the difference in relationship between trait values and fitness between ancestral and descendant populations. Observed differences between traits and fitness between sampling time points do not reflect changes in selection patterns *per se*, but more accurately indicate differences in how populations respond to a shared selective regime. These differences could be due to correlated traits or differing trait distributions of the ancestral or descendant populations. We controlled for different trait distributions by standardizing trait measurements across sampling years to hold distribution constant. Additionally, as a conceptual check prior to standardization, we directly contrast trait distributions between sampling years. We found no shifts in the trait distribution for floral traits, but did observe a lower leaf count distribution in descendant populations (Fig. S4). Given that there was no corresponding correlational selection or change in covariance with leaf count, any observed changes in the fitness-leaf count relationship are unlikely to be due to constraints. The stable distribution in flowering phenology between sampling years supports the idea that a change in response to the selective environment is indicative of a decline in adaptive potential.

Traits are again standardized such that trait variance equals one. We report differentials and gradients so we can assess both the total (indirect and direct) selective effects on a trait, as well as isolate which traits are directly under selection, and are thus causal drivers of evolution in this set of pollination-related traits (Tables S2, S3).

We estimated selection differentials ($S$) using a regression of relative fitness on each trait separately in a model containing a linear term for the trait of interest and controlling for block as a random effect:

$$\text{Relative Fitness} \sim \text{Trait} + \text{Block}$$

We further looked at total balancing or disruptive selection using a similar model with linear and quadratic terms for the trait of interest:

$$\text{Relative Fitness} \sim \text{Trait} + \text{Trait}^{\wedge}2 + \text{Block}$$

Selection gradients were calculated using the multivariate regression model described above and significance of effect on fitness is assessed using a type III ANOVA.

**Pollination frequency and SEM model selection**–For each plant, we sum all observed approaches and visits to obtain individual approach and visit frequencies. Since the length of observation period and number of rounds of observation were held constant across all plots, this represents a time-standardized metric of insect pollinator visitation to each plant. For corolla width, corolla length, ASD, and nectar sucrose content, we use the mean of four randomly selected flowers distributed over the growing season as the trait value for each plant, whereas date of first flower and plant size are single values represented as Julian date and leaf count, respectively. Residuals from a linear regression of plot on trait to remove were used in all further analyses to remove plot effects.

To determine whether changes in trait values over time are mediated by pollinator behavior, we use structural equation modeling (SEM) to evaluate seven *a priori* hypotheses exploring the relationship between phenotype, pollinator visitation, and fitness (Table S4). Structural equation modeling assesses causal relationships between variables, both direct and indirect, by taking two inputs: 1) qualitative causal assumptions (*i.e.* some a priori biological hypothesis about how parameters interact), and 2) empirical data, to then derive two logical conclusions: a statistical measure of fit for the model that describes the implications of the assumptions (*i.e.* do the assumptions about causality between parameters adequately describe the covariance structure of the data), and coefficients representing the strength and significance of causal relationships between parameters (Bollen & Pearl, 2012). Even if a model contains significant coefficients, poor model fit casts doubt on the assumptions included in the model structure. An accepted model does not prove causal assumptions (*i.e.* model-reality consistency); however it does indicate higher plausibility of estimated relationships by demonstrating model-data consistency through variance-covariance structure. As such, we first assess model fit for each of our seven models that represent hypotheses about how plant traits influence fitness either directly or indirectly *via* pollinators (Table S4).

These hypotheses include five nested models A-E (*i.e.* removal and inclusion of directional relationships with no change in the causal order of relationships), and two additional non-nested models, F and G (Table S4, S5). Models A-C test the degree to which the impact of floral traits on fitness is mediated through pollinators. Model A proposes that the fitness effect of all traits other than plant size are fully mediated through pollinator interactions. In turn, model B proposes that the fitness impact of floral traits is only partially mediated by pollinators, and model C proposes that the impact of floral traits on fitness is not mediated through pollinators at all. Removing the causal link between plant size and fitness tests the assumption that fitness is limited by internal resources (Model D) and removing direct links from functional traits to pollinator foraging tests whether all pollinator foraging choice occurs prior to entering the corolla such that fitness impacts are mediated by signaling cues that can be perceived from a distance (Model E). Finally, we test two non-nested models: Model F proposes that variation in traits is all mediated through plant size, adding a causal relationship between leaf count and all other traits, and Model G assesses the possibility that relationships between traits and pollinator behavior are correlated, but not causal (*i.e.* a bi-directional rather than uni-directional relationship).

We use piecewiseSEM in R, which allows for more flexible data assumptions, namely that the data is not required to be multivariate normal, and generalized linear models can be fit for non-Gaussian data types such as count (Lefcheck, 2016). piecewiseSEM implements a log-likelihood based goodness of fit measure that produces a $\chi^2$ statistic comparing model-implied variance-covariance relationships with actual variance-covariance relationships in the data. A significant p-value ($< 0.05$) in this case indicates a significant difference between the relationships proposed by the model and those present in the data, meaning the model is not an appropriate fit to the data. Conversely, a non-significant $\chi^2$ comparison indicates that the model does fit the data. In one case (model B), the model does not have any degrees of freedom, so we use Akaike's information criterion (AIC) to compare it to a similar model assuming plant size influences only pollinator approach frequency, not pollinator foraging frequency, as it is unlikely that plant size influences pollinator foraging choice after the pollinator has already approached the plant. We find that the AIC is significantly (>2 AIC units) lower for this second model, so proceed with model selection using a modified model B that does not include plant size as an effect on pollinator foraging frequency (Table S5).

**Causal relationships and power analysis**–The only model with a reliable fit to the observed data was Model B, which tested the hypothesis that the effect of floral traits on fitness is partially mediated through pollinator behavior (Model B, $\chi^2 = 0.246$, $p = 0.619$, Table S5). All other models failed to explain the correlation matrix of the data ($p < 0.05$). For Model B, we perform multigroup analysis in piecewiseSEM to calculate direct and indirect effects and calculate path coefficients for ancestral populations. Direct effects represent standardized partial regression coefficients, and indirect effects of traits are calculated by multiplying direct effect coefficients along any given path from trait to pollinator behavior to plant fitness. Finally, we use pwrSEM (Wang & Rhemtulla, 2021) to calculate the statistical power of all our models and determine whether the lack of a significant path between a functional trait, pollinator behavior, and fitness is a true lack of relationship, not a lack of power to detect a relationship. Power is calculated as the percent of 5,000 simulated samples with a parameter estimate not significantly different from the coefficient estimated from the true data (Table S7).

**Phenotypic Evolution**–To assess whether trait values change in alignment with our expectations given the pattern of selection and genetic covariances, we test for realized phenotypic evolution between ancestral and descendant populations. To do this, we first perform a linear mixed model using the lme4 package (Bates *et al.*, 2015) in R with year, population, and the interaction of year and population as fixed effects and block as a random effect to control for spatial differences across the common garden. All phenotypic traits were log-transformed to adhere to assumptions of residual normality. Each trait was analyzed in a separate model of the following general form:

$$\text{Trait} \sim \text{Year*Population} + \text{Block}$$

We assessed the significance of differences in trait values between sampling years and whether responses differed between sampling populations, using a posthoc analysis of estimated marginal means for each trait in each year. To do so, we used the emmeans package in R (Lenth, 2025) and performed a t-ratio test comparing ancestral to descendant populations. Flowering phenology displayed a repeatable bimodal distribution across both cohorts, consistent with previous resurrection and greenhouse studies of these populations (Bishop et al. 2023).

### Figures

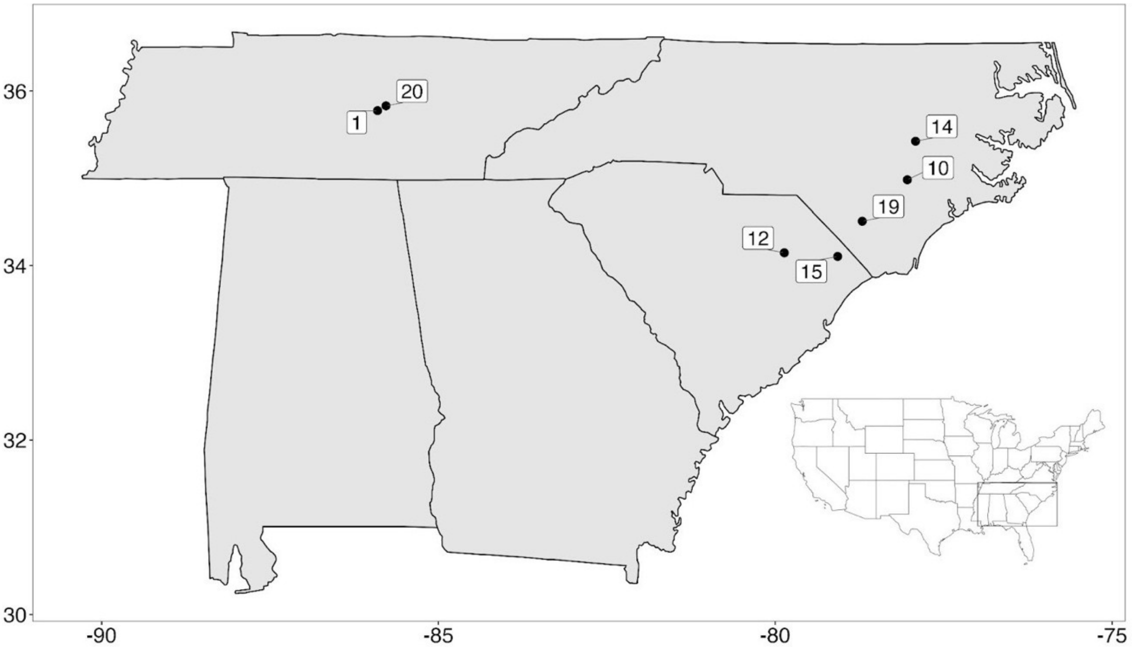


**Fig. S1.** Distribution of seven sampling localities of *I. purpurea* labeled with population number. All populations were sampled from the edge of agricultural soy and maize fields.

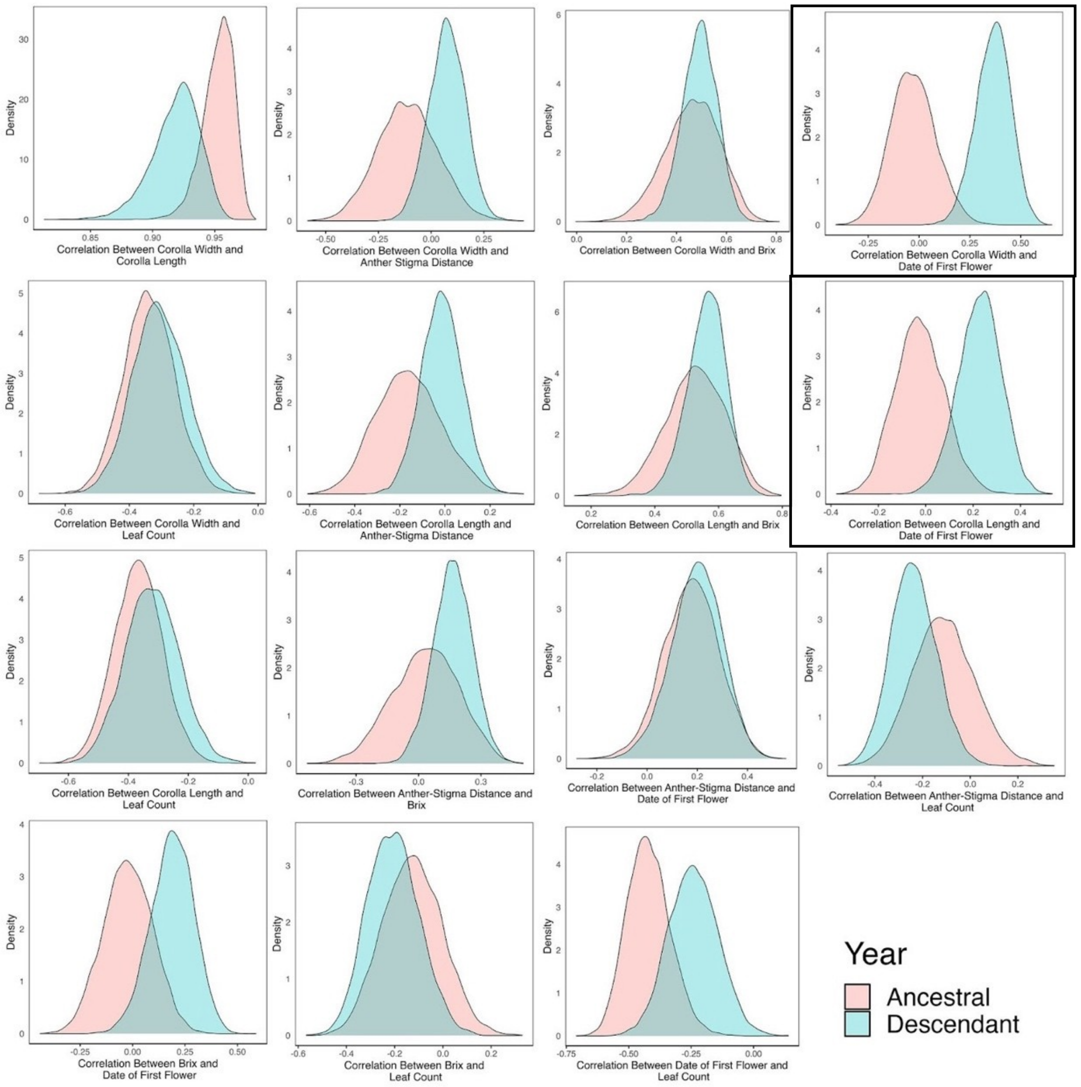


**Fig. S2.** Comparison of the distribution of correlations for each pairwise trait combination in ancestral and descendant populations. Each distribution is attained by bootstrap resampling trait values 10,000 times. The distribution of the Pearson's correlation coefficient of all samples is plotted with trait pairs that show a significant difference between ancestral and descendant populations outlined with a black box. Significance is determined by the difference between the ancestral and descendant coefficient having the same sign for 95% or more re-sampled populations.

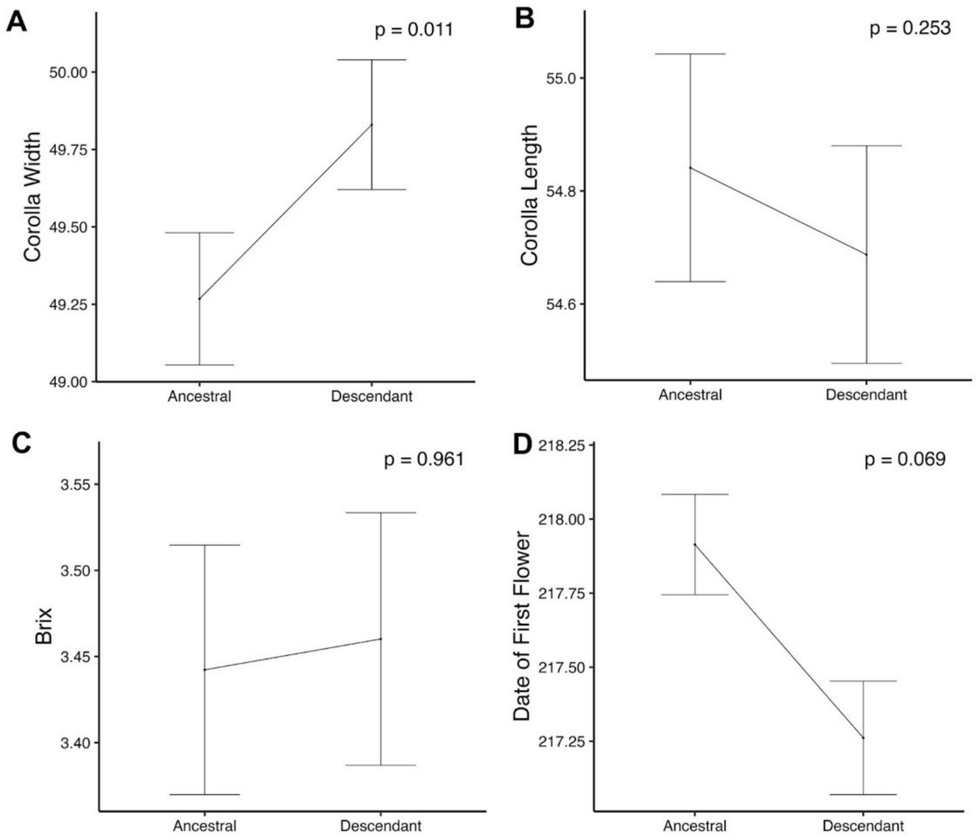


**Fig. S3.** Estimated marginal means plotted with standard deviation for phenotypic traits that show significant correlational selection. Traits are plotted with raw values to display a biologically interpretable degree of change, but significance is assessed using log-transformed traits analyzed with a linear mixed model of the general form Trait ~ Year*Population + Block where block is the only random effect. Reported p-values come from a post hoc t-ratio test comparing estimated marginal means between the years.

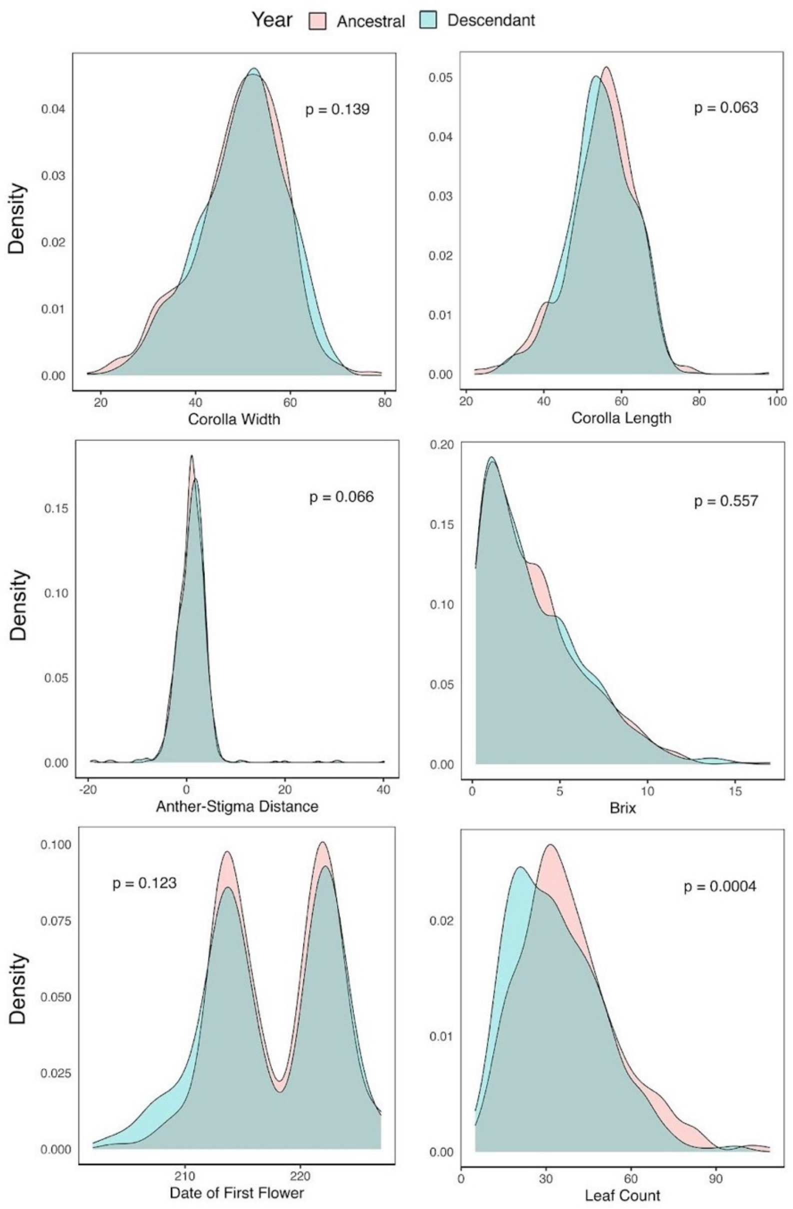


**Fig. S4.** Distribution of trait values across all populations in ancestral and descendant populations for corolla width, corolla length, anther-stigma distance, nectar sucrose (°Brix), date of first flower, and leaf count. Significant differences in trait distribution between years are displayed as p-values from separate Kolmogorov–Smirnov tests for each trait.

### Tables

**Table S1.** Phenotypic change from ancestral to descendant populations using estimated marginal means. Effect sizes (mean and standard error) are shown with non-transformed data such that the scale of change is maintained, but significance is assessed using trait data standardized to a mean of zero and a standard deviation of one. Trait change is evaluated using a linear mixed model of the general form Trait ~ Year*Population + Block where block is the only random effect. Reported t-ratio, degrees of freedom, and p-values come from a post hoc t-ratio test comparing estimated marginal means between the years.

| | | Corolla Width (mm) | Corolla Length (mm) | ASD (mm) | Nectar Sucrose (°Brix) | Flowering Phenology (Julian Date) | Plant Size (Leaf Count) |
|---|---|---|---|---|---|---|---|
| Mean | Ancestral | 47.2 | 52.7 | 0.44 | 3 | 218 | 37.2 |
| | Descendant | 48.1 | 53 | 0.59 | 3 | 217.3 | 33.0 |
| Standard Error | Ancestral | 3.03 | 3.17 | 0.73 | 0.61 | 0.49 | 3.69 |
| | Descendant | 3.02 | 3.17 | 0.73 | 0.61 | 0.48 | 3.68 |
| t-ratio | | -2.53 | -0.85 | -1.08 | -0.02 | 1.78 | 3.67 |
| Degrees Freedom | | 1888 | 1885 | 1860 | 1432 | 799 | 799 |
| p-value | | 0.01 | 0.40 | 0.28 | 0.98 | 0.07 | 3.0e-4 |

**Table S2.** Total linear and nonlinear selection (S) on floral traits (corolla width, corolla length, ASD, nectar sucrose (°Brix), and date of first flower) as well as plant size in ancestral and descendant populations. Selection differential coefficients, standard errors, and p-values for each trait are evaluated using a separate linear mixed model for each trait of the form Relative Fitness ~ Year*(Trait + Trait^2) + Block with block as a random effect and quadratic coefficients and standard errors multiplied by two. F-values for the year by trait interaction from an ANCOVA are shown with a significant effect indicating a difference in total selection on the trait between years ($p < 0.1$~, $p < 0.05$*, $p < 0.01$**, and $p < 0.001$***).

| | | **Ancestral** | | | **Descendant** | | | **F-values from ANCOVA** |
|---|---|---|---|---|---|---|---|---|
| **Trait** | | **S** (linear) or **γ** (quadratic) | **SE** | **p-value** | **S** (linear) or **γ** (quadratic) | **SE** | **p-value** | |
| **Corolla Width** | Linear | 0.14 | 0.04 | 1.5e-4 | 0.14 | 0.03 | <0.0001 | 0.01 |
| | Quadratic | -0.12 | 0.05 | 0.030 | 0.02 | 0.05 | 0.619 | 3.75* |
| **Corolla Length** | Linear | 0.11 | 0.04 | 0.002 | 0.06 | 0.03 | 0.031 | 0.80 |
| | Quadratic | -0.05 | 0.05 | 0.328 | -0.05 | 0.04 | 0.218 | 0.01 |
| **ASD** | Linear | 0.03 | 0.04 | 0.466 | 0.007 | 0.03 | 0.788 | 0.35 |
| | Quadratic | 3e-03 | 0.03 | 0.907 | -0.02 | 0.01 | 0.070 | 0.32 |
| **Nectar Sucrose (°Brix)** | Linear | 0.04 | 0.04 | 0.322 | 0.07 | 0.03 | 0.04 | 0.03 |
| | Quadratic | 0.14 | 0.06 | 0.022 | 0.01 | 0.05 | 0.902 | 3.40~ |
| **Date of First Flower** | Linear | -0.20 | 0.08 | 0.015 | 0.04 | 0.05 | 0.511 | 5.44* |
| | Quadratic | -0.02 | 0.17 | 0.925 | -0.24 | 0.08 | 0.002 | 1.22 |
| **Leaf Count** | Linear | 0.06 | 0.07 | 0.343 | -0.04 | 0.06 | 0.503 | 0.71 |
| | Quadratic | -0.04 | 0.09 | 0.621 | -0.08 | 0.09 | 0.384 | 0.07 |

**Table S3.** Direct selection (**β**) acting on floral traits: corolla width, corolla length, ASD, Nectar Sucrose (°Brix), and date of first flower in ancestral and descendant populations. Since plant size is included as a control for indirect selection, not a primary effect, non-significant leaf count terms are removed from the model. Selection coefficients, standard errors, and p-values for each trait are obtained using a linear mixed model with relative fitness regressed on trait with linear, doubled quadratic, and cross-product terms. F-values for the year by trait interaction from an ANCOVA are shown with a significant effect indicating a difference in total selection on the trait between years ($p < 0.1$~, $p < 0.05$*, $p < 0.01$**, and $p < 0.001$***).

| | | **Ancestral** | | | **Descendant** | | | **F-values from ANCOVA** |
|---|---|---|---|---|---|---|---|---|
| **Trait** | | **β** | **SE** | **p-value** | **β** | **SE** | **p-value** | |
| **Corolla Width** | Linear | 0.34 | 0.25 | 0.175 | 0.45 | 0.16 | 0.007 | 0.63 |
| | Quadratic | -0.37 | 0.45 | 0.405 | 0.45 | 0.26 | 0.090 | 0.53 |
| **Corolla Length** | Linear | -0.12 | 0.25 | 0.625 | -0.37 | 0.17 | 0.030 | 2.22 |
| | Quadratic | -0.35 | 0.45 | 0.441 | -0.22 | 0.24 | 0.353 | 1.92 |
| **ASD** | Linear | 0.07 | 0.11 | 0.515 | -0.002 | 0.06 | 0.968 | 0.48 |
| | Quadratic | -0.06 | 0.08 | 0.413 | -0.04 | 0.04 | 0.322 | 0.40 |
| **Nectar Sucrose (°Brix)** | Linear | -0.15 | 0.13 | 0.235 | -0.01 | 0.09 | 0.877 | 0.01 |
| | Quadratic | 0.22 | 0.08 | 0.013 | 0.10 | 0.06 | 0.091 | 1.52 |
| **Date of First Flower** | Linear | -0.28 | 0.13 | 0.036 | -0.05 | 0.06 | 0.393 | 8.96** |
| | Quadratic | 0.09 | 0.15 | 0.509 | -0.13 | 0.04 | 0.005 | 3.62~ |

**Table S4.** Seven hypothesized models of the direct and indirect effects that determine pollinator visitation frequency and plant fitness. Models A – E are nested models testing the degree to which trait-based effects on fitness are moderated through pollinator behavior. Models F and G are non-nested and test whether trait effects are reducible to plant size or are equally well explained by correlative rather than causative relationships. All models include bi-directional paths accounting for correlations between functional traits.

| **Model** | **Path Alterations** | **Biological Hypothesis** |
|---|---|---|
| A | No direct paths from traits (except plant size) to fitness | Effect of floral traits on fitness is entirely mediated through pollinator behavior, fitness is limited by internal resources |
| B | Indirect and direct paths from traits to fitness | Effect of floral traits on fitness is partially mediated through pollinator behavior |
| C | No indirect paths from traits to fitness | Effect of floral traits on fitness is not mediated through pollinator behavior at all |
| D | No causal link between plant size and fitness | Fitness is not limited by internal resources |
| E | No paths from traits to pollinator foraging | All pollinator foraging choice occurs prior to entering corolla such that fitness impacts are mediated by signaling cues that can be perceived from a distance |
| F | Additional path between leaf count and all other traits | Variation in functional traits is mediated through plant size |
| G | Bi-directional instead of uni-directional paths between traits and pollinator visitation/foraging | Relationships between traits and pollinator behavior are correlated, not causal |

**Table S5.** A comparison of goodness of fit for alternative path diagrams (Figure 3) using structural equation modeling. Nonsignificant $\chi^2$ values suggest that the models do not deviate significantly from the observed data. AIC is Akaike's information criterion and df is degrees of freedom in the model. Models that show nonsignificant $\chi^2$ and minimize AIC provide the most reliable fit to the observed data.

| **Model** | **Dataset** | $\chi^2$ | **df** | **p-val** | **AIC** |
|---|---|---|---|---|---|
| A | Full | 15.390 | 6 | 0.017 | 3065 |
| B | Full | 0.236 | 1 | 0.627 | 3060 |
| C | Full | 23.465 | 6 | 0.001 | 2206 |
| D | Full | 13.074 | 2 | 0.001 | 3070 |
| E | Full | 23.465 | 6 | 0.001 | 3073 |
| F | Full | 388.58 | 11 | 0 | 10281 |
| G | Full | 480.52 | 15 | 0 | 12638 |

**Table S6.** Direct, indirect, and total effects of floral characteristics and pollinator approach and visit frequency on plant fitness for Model B (Figure 4). Direct effects (DE) are standardized partial regression coefficients from the SEM. Indirect effects (IE) are the product of all direct effects along a given path. If multiple indirect paths between trait and fitness exist, coefficients of the independent paths are summed to get a total indirect effect on fitness. Total effects (TE) are the sum of indirect and direct effects, and non-significant paths are marked with NA.

| Trait | Year | **Approach** | **Visit** | **Fitness** | | |
|---|---|---|---|---|---|---|
| | | DE | DE | DE | IE | TE |
| **Corolla Width** | 2003 | 0.250 | 0.337 | NA | 0.074 | 0.074 |
| | 2012 | 0.274 | 0.349 | 0.428 | NA | 0.428 |
| **Corolla Length** | 2003 | NA | NA | NA | NA | NA |
| | 2012 | NA | NA | -0.825 | NA | -0.825 |
| **ASD** | 2003 | NA | 0.109 | NA | 0.019 | 0.019 |
| | 2012 | NA | 0.207 | NA | NA | NA |
| **Nectar Sucrose (°Brix)** | 2003 | 0.148 | NA | NA | 0.009 | 0.009 |
| | 2012 | 0.216 | NA | NA | NA | NA |
| **Date of First Flower** | 2003 | NA | NA | -0.325 | NA | -0.325 |
| | 2012 | NA | NA | NA | NA | NA |
| **Leaf Count** | 2003 | NA | NA | 0.259 | NA | 0.259 |
| | 2012 | NA | NA | 0.212 | NA | 0.212 |
| **Approach** | 2003 | NA | 0.373 | NA | 0.064 | 0.064 |
| | 2012 | NA | 0.353 | NA | NA | NA |
| **Visit** | 2003 | NA | NA | 0.171 | NA | 0.171 |
| | 2012 | NA | NA | NA | NA | NA |

**Table S7.** Power analysis of Model B. 'Coefficient' is the estimated coefficient from the SEM, 'Median Coefficient' is the median coefficient estimate of 5000 simulations, and 'Power' is the percent of 5000 simulations with a parameter estimate not significantly different than the coefficient estimated from the true data ($p < 0.05$) as determined by pwrSEM (*46*). Yellow highlight indicates where lower power and lack of a significant signal coincide, indicating that there may be a significant relationship, but a lack of power to detect it.

| | **2003** | | | **2012** | | |
|---|---|---|---|---|---|---|
| **Parameter** | **Coefficient** | **Median Coefficient** | **Power** | **Coefficient** | **Median Coefficient** | **Power** |
| relFit ~ Visit | 0.17 | 0.17 | 1 | -0.12 | -0.12 | 0.99 |
| relFit ~ Approach | 0.07 | 0.07 | 0.16 | 0.08 | 0.08 | 0.18 |
| relFit ~ CWresid | -0.3 | -0.3 | 1 | 0.45 | 0.45 | 1 |
| relFit ~ CLresid | 0.19 | 0.19 | 1 | -0.8 | -0.8 | 1 |
| relFit ~ ASDresid | 0.06 | 0.05 | 0.13 | 0.08 | 0.08 | 0.31 |
| relFit ~ Bresid | 0.07 | 0.07 | 0.19 | 0.11 | 0.11 | 0.63 |
| relFit ~ FFresid | -0.32 | -0.32 | 1 | 0.13 | 0.13 | 1 |
| relFit ~ LCresid | 0.26 | 0.26 | 1 | 0.21 | 0.21 | 1 |
| Approach ~ CWresid | 0.24 | 0.24 | 1 | 0.26 | 0.26 | 1 |
| Approach ~ CLresid | -0.06 | -0.06 | 0.81 | -0.06 | -0.06 | 0.71 |
| Approach ~ ASDresid | 0.04 | 0.04 | 0.09 | 0.05 | 0.05 | 0.16 |
| Approach ~ Bresid | 0.15 | 0.15 | 0.58 | 0.22 | 0.22 | 0.99 |
| Approach ~ FFresid | 0.2 | 0.2 | 1 | -0.14 | -0.14 | 1 |
| Approach ~ LCresid | 0.12 | 0.12 | 1 | 0.09 | 0.09 | 1 |
| Visit ~ CWresid | 0.29 | 0.29 | 1 | 0.3 | 0.3 | 0.98 |
| Visit ~ CLresid | -0.09 | -0.09 | 0.38 | -0.08 | -0.08 | 0.26 |
| Visit ~ ASDresid | 0.21 | 0.22 | 0.2 | 0.27 | 0.27 | 0.38 |
| Visit ~ Bresid | 0.01 | 0 | 0.06 | 0.01 | 0.01 | 0.06 |
| Visit ~ FFresid | 0.21 | 0.21 | 0.93 | -0.12 | -0.12 | 0.54 |
| Visit ~ Approach | 0.37 | 0.36 | 0.87 | 0.35 | 0.34 | 0.52 |

**Table S8.** For each trait, the mean and standard error for $\mathbf{\Delta z_i}$ and $\mathbf{\Delta z_{i\_nc}}$, including population 12. $\Delta z$ is calculated with standardized traits such that variance(maternal line) = 1. Ancestral and descendent populations are noted as A or D, respectively. ***R*** estimates for ancestral and descendant populations including population 12 are displayed at bottom, showing a similar, but less extreme, trend as results excluding population 12 with descendant populations displaying greater constraint than ancestral.

| | **Year** | $\Delta \bar{z}$ | **SE** | $\Delta \bar{z}_{nc}$ | **SE** |
|---|---|---|---|---|---|
| **Corolla Width** (mm) | A | -3.14E-5 | 6.92E-4 | 0.13 | 7.88E-4 |
| | D | 0.02 | 2.58E-04 | 0.08 | 5.88E-04 |
| **Corolla Length** (mm) | A | -5.95E-3 | 6.58E-4 | -0.08 | 5.05E-4 |
| | D | 0.01 | 2.14E-04 | -0.05 | 4.08E-04 |
| **ASD** (mm) | A | -0.04 | 7.02E-04 | 2.49E-03 | 1.85E-05 |
| | D | -0.02 | 3.51E-04 | -0.02 | 1.71E-04 |
| **Nectar Sucrose** (°Brix) | A | -0.01 | 6.94E-04 | -0.03 | 3.23E-04 |
| | D | 0.01 | 2.22E-04 | -1.43E-03 | 1.33E-05 |
| **Flowering Phenology** (Julian Date) | A | -0.12 | 7.00E-04 | -0.07 | 3.18E-04 |
| | D | -0.01 | 3.10E-04 | -0.03 | 8.68E-05 |
| **Plant Size** (Leaf Count) | A | 0.16 | 1.16E-03 | 0.14 | 1.03E-03 |
| | D | 0.02 | 2.90E-04 | 0.02 | 1.41E-04 |
| $\boldsymbol{R}_{ancestral}$ = 0.56 | | | $\boldsymbol{R}_{descendant}$ = 0.11 | | |

**Table S9.** By-population analysis of $\Delta z_i$ and $\Delta z_{i\_nc}$. All trait measurements are scaled and standardized to a mean of zero and standard deviation of one. For each year within each population, 2,000 bootstrap samples are used to calculate mean and standard error of $\Delta z_i$ and $\Delta z_{i\_nc}$. Significance is defined as the percent of bootstrap samples with a threshold of 95% where $\Delta z_i - \Delta z_{i\_nc}$ is either above or below zero, depending on the expected difference calculated from the actual data. ***R*** cannot be calculated for individual populations due to insufficient power for interaction terms in the selection gradient model.

| Pop | Year | Trait | $\Delta z_i$ | SE | $\Delta z_{i\_nc}$ | SE |
|---|---|---|---|---|---|---|
| 1 | Ancestral | Corolla Width | -1.07 | 0.04 | 5.33 | 0.06 |
| | | Corolla Length | -1.37 | 0.04 | -8.12 | 0.08 |
| | | ASD | 0.01 | -0.68 | 4.5E-03 | 0 |
| | | Brix | -0.28 | 0.02 | -0.73 | 0.01 |
| | | Date of First Flower | -0.97 | 0.02 | 0.19 | 0.01 |
| | | Leaf Count | 0.29 | 0.01 | 0.40 | 2.7E-03 |
| | Descendant | Corolla Width | -0.10 | 4.40E-03 | 0.78 | 0.01 |
| | | Corolla Length | -0.13 | 0.01 | -0.75 | 0.01 |
| | | ASD | 0.01 | 0.01 | -0.33 | 3.90E-03 |
| | | Brix | -0.03 | 2.80E-03 | -0.19 | 1.80E-03 |
| | | Date of First Flower | -0.04 | 4.50E-03 | 0.01 | 3.90E-03 |
| | | Leaf Count | 0.11 | 0.01 | 0.02 | 0.01 |
| 10 | Ancestral | Corolla Width | 0.94 | 0.02 | 13.2 | 0.26 |
| | | Corolla Length | 0.49 | 0.02 | -12 | 0.23 |
| | | ASD | -1.53 | 0.02 | -1.22 | 0.02 |
| | | Brix | -0.72 | 0.02 | -1.32 | 0.03 |
| | | Date of First Flower | -0.33 | 0.01 | 0.73 | 0.02 |
| | | Leaf Count | -0.28 | 0.02 | -0.54 | 0.02 |
| | Descendant | Corolla Width | 0.53 | 0.01 | 0.11 | 0.02 |
| | | Corolla Length | 0.2 | 0.01 | 0.35 | 0.01 |
| | | ASD | 2 | 0.03 | 0.44 | 0.01 |
| | | Brix | -1.02 | 0.02 | -0.17 | 3.70E-03 |
| | | Date of First Flower | 0.95 | 0.02 | 0.84 | 0.02 |
| | | Leaf Count | -1.04 | 0.02 | -0.18 | 4.00E-03 |
| 12 | Ancestral | Corolla Width | 0.19 | 0.01 | 1.09 | 0.02 |
| | | Corolla Length | 0.17 | 0.01 | -0.37 | 0.02 |
| | | ASD | 0.07 | 0.01 | -0.35 | 0.01 |
| | | Brix | 0.01 | 0.01 | -0.62 | 0.01 |
| | | Date of First | 0.07 | 0.01 | 0.44 | 0.01 |

| | | | | | | |
|---|---|---|---|---|---|---|
| | | Flower | | | | |
| | | Leaf Count | 0.02 | 0.01 | 1 | 0.02 |
| | Descendant | Corolla Width | 0.08 | 0.01 | 0.84 | 0.01 |
| | | Corolla Length | 0.38 | 0.01 | -0.82 | 0.01 |
| | | ASD | -0.16 | 0.01 | -1.39 | 0.02 |
| | | Brix | 0.41 | 0.01 | 1 | 0.01 |
| | | Date of First Flower | 0.33 | 3.70E-03 | 0.7 | 0.01 |
| | | Leaf Count | 0.18 | 0.01 | 0.93 | 0.02 |
| 14 | Ancestral | Corolla Width | -1.7 | 0.16 | 4.01 | 0.18 |
| | | Corolla Length | -2.11 | 0.19 | -4.26 | 0.29 |
| | | ASD | -0.01 | 0.08 | 0.28 | 0.09 |
| | | Brix | 0.05 | 0.07 | 1.74 | 0.1 |
| | | Date of First Flower | -3.67 | 0.22 | 0.31 | 0.04 |
| | | Leaf Count | 4.88 | 0.26 | 4.42 | 0.26 |
| | Descendant | Corolla Width | -0.97 | 0.03 | 4.79 | 0.07 |
| | | Corolla Length | -1.69 | 0.04 | -6.17 | 0.1 |
| | | ASD | -1.17 | 0.04 | 1.07 | 0.05 |
| | | Brix | -1.12 | 0.03 | -1.08 | 0.02 |
| | | Date of First Flower | 1.03 | 0.02 | -0.71 | 0.02 |
| | | Leaf Count | -0.56 | 0.02 | -1.03 | 0.02 |
| 15 | Ancestral | Corolla Width | -0.43 | 0.01 | 0.58 | 0.01 |
| | | Corolla Length | 0.02 | 0.02 | -0.52 | 0.02 |
| | | ASD | -0.87 | 0.03 | 0.76 | 0.01 |
| | | Brix | 2.59 | 0.12 | 1.86 | 0.04 |
| | | Date of First Flower | -0.33 | 0.01 | 0.13 | 3.10E-03 |
| | | Leaf Count | 0.14 | 4.10E-03 | 0.04 | 2.20E-03 |
| | Descendant | Corolla Width | 0.1 | 0.02 | -1.83 | 0.05 |
| | | Corolla Length | 0.14 | 0.01 | 1.51 | 0.06 |
| | | ASD | 0.54 | 0.02 | 1.78 | 0.03 |
| | | Brix | -0.07 | 0.01 | 0.16 | 0.01 |
| | | Date of First Flower | -0.56 | 0.02 | -1.57 | 0.02 |
| | | Leaf Count | 0.2 | 0.01 | -0.23 | 0.01 |
| 19 | Ancestral | Corolla Width | -2.85 | 0.08 | 3.36 | 0.08 |
| | | Corolla Length | -4.53 | 0.12 | -3.55 | 0.11 |
| | | ASD | -6.67 | 0.15 | -1.21 | 0.04 |
| | | Brix | -2.48 | 0.07 | 0.18 | 0.01 |
| | | Date of First Flower | -2.47 | 0.04 | -2.13 | 0.03 |

| | | | | | | |
|---|---|---|---|---|---|---|
| | | Leaf Count | 8.46 | 0.18 | 0.91 | 0.01 |
| | Descendant | Corolla Width | 0.69 | 0.04 | 2.72 | 0.07 |
| | | Corolla Length | 0.5 | 0.03 | 1.47 | 0.04 |
| | | ASD | 0.35 | 0.03 | 1.15 | 0.02 |
| | | Brix | 0.02 | 0.02 | -2.09 | 0.04 |
| | | Date of First Flower | -0.08 | 0.01 | -0.66 | 0.02 |
| | | Leaf Count | 1.64 | 0.04 | 1.5 | 0.05 |
| 20 | Ancestral | Corolla Width | 1.18 | 0.02 | 3.16 | 0.04 |
| | | Corolla Length | 0.45 | 0.01 | -2.23 | 0.02 |
| | | ASD | 0.73 | 0.01 | 0.1 | 0.01 |
| | | Brix | 0.12 | 0.01 | -0.72 | 0.02 |
| | | Date of First Flower | 1.43 | 0.01 | 1.31 | 0.02 |
| | | Leaf Count | -0.32 | 0.01 | 1.46 | 0.03 |
| | Descendant | Corolla Width | 4.68 | 0.04 | -8.07 | 0.13 |
| | | Corolla Length | 4.11 | 0.04 | 9.88 | 0.14 |
| | | ASD | 3.19 | 0.04 | 0.77 | 0.02 |
| | | Brix | 3.02 | 0.03 | 0.27 | 4.50E-03 |
| | | Date of First Flower | 2.78 | 0.03 | 1.86 | 0.03 |
| | | Leaf Count | -1.77 | 0.02 | 0.3 | 0.02 |
